# Read mismapping from segmental duplications drives spurious *trans* associations

**DOI:** 10.1101/2025.10.08.681258

**Authors:** Alber Aqil, Omer Gokcumen

## Abstract

Heritable variation in gene regulation shapes both disease risk and evolution. *Trans*-acting quantitative trait loci (*trans*-QTLs) are the primary source of this variation within species, but they remain difficult to identify correctly due to false positives stemming from RNA and DNA read mismapping. Here, we identify four categories of RNA and DNA mismapping due to segmental duplications that lead to false-positive *trans*-QTLs. Such mismapping leaves a footprint in the form of long-range genotypic disequilibrium, which we use to identify spurious *trans*-QTLs. Applying this method to Genotype-Tissue Expression (GTEx) *trans*-QTL, we conservatively estimate that 14% of expression QTLs, 10% of splicing QTLs, and 3% of protein QTLs are artifacts. More broadly, this framework could be applied to flag spurious distant peaks in genome-wide association studies, Hi-C contacts, and co-expression between distant genes.

## Introduction

Heritable variation in gene regulation underlies both disease risk (Aqil et al. 2025) and rapid phenotypic evolution (King and Wilson 1975; Singh et al. 2025). Large-scale undertakings, such as the Genotype-Tissue Expression (GTEx) project, have identified specific genetic variants within Quantitative Trait Loci (QTL) that lead to heritable variation in gene regulation. Such regulatory variants can act either in *cis* or *trans* with respect to a given gene. On the one hand, *cis*-regulatory variants (henceforth, *cis-*QTLs) tend to be located close to the target gene and affect allele-specific regulatory regions such as promoters and enhancers. On the other hand, *trans*-regulatory variants (henceforth, *trans*-QTLs) can be located anywhere in the genome, and affect the target gene’s expression from both alleles primarily via a diffusible molecule like RNA or protein. For example, variants in genes for transcription factors (Võsa et al. 2021), splicing factors (Verta and Jacobs 2022), and epigenetic regulators (Tulstrup et al. 2021) can alter expression levels, isoform usage, and methylation patterns of many distant target genes.

Previous studies suggest that *trans*-QTLs are the primary source of gene-expression variation within species (Emerson et al. 2010; Metzger et al. 2017; Liu et al. 2019; Gokhman et al. 2021). Yet, correctly identifying these variants remains challenging due to three reasons. Firstly, *trans*-QTLs segregating at frequencies required by QTL detection pipelines have small effect sizes. This is because even when a *trans*-QTL and a *cis*-QTL for a target gene have the same effect size, the *trans*-QTL is more deleterious because of its additional effects on other genes (Vande Zande et al. 2022; Vande Zande and Wittkopp 2022; Kimura and Ohta 1974). Consequently, *trans*-QTLs undergo intense purifying selection, such that those that persist at appreciable frequencies tend to have smaller effects than their *cis* counterparts (Fu et al. 2013; Mähler et al. 2017), making their identification difficult. This leads to a false-negative problem. Secondly, this false-negative problem is exacerbated by the stringent multiple-hypothesis testing required for testing millions of gene-variant pairs for trans-QTL associations. This results in a stricter significance threshold, which further reduces the power to detect the small-effect *trans*-QTLs. Thirdly, reads from a *cis*-regulatory region or its target gene can map to a paralog elsewhere in the genome. In such cases, *cis*-QTLs, which have larger effect sizes, may masquerade as *trans*-QTLs for a paralogous gene, leading to a high rate of false positives. Taken together, the false negatives due to small effects of genuine *trans*-QTLs and the heavy burden of multiple testing, and false positives due to read mismapping, mean that many of the apparent *trans*-QTLs detected are likely artifacts.

Given the false-negative problem, most genuine *trans*-QTLs, which have small effect sizes, are not detected. The only genuine *trans*-QTLs that are detected should be outliers in terms of having large effect sizes. Such variants are of particular evolutionary and mechanistic interest. In particular, they may be maintained by balancing selection (broadly construed) (Aqil et al. 2023), undergoing positive selection, or exerting gene-specific effects despite acting through a diffusible molecule. Identifying these rare cases requires distinguishing genuine *trans*-QTLs from spurious *trans*-QTLs resulting from the mismapping of RNA and DNA reads. This report provides a framework for making this distinction.

## Results

We identify four major categories of read mismapping that can cause *cis*-QTLs to appear as *trans*-QTLs. Categories 1–2 involve duplications present in the reference genome, affecting either the gene (**Figure 1A**) or its *cis*-regulatory element (**Figure 1B**). Categories 3–4 involve polymorphic duplications absent from the reference, again affecting either the gene (**Figure 1C**) or its *cis*-regulatory element (**Figure 1D**). We describe these categories of mismapping in detail in the Figure 1 legend for expression QTLs (eQTLs), but the same principles apply to splicing, methylation, and protein QTLs in different combinations.

**Figure 1.**
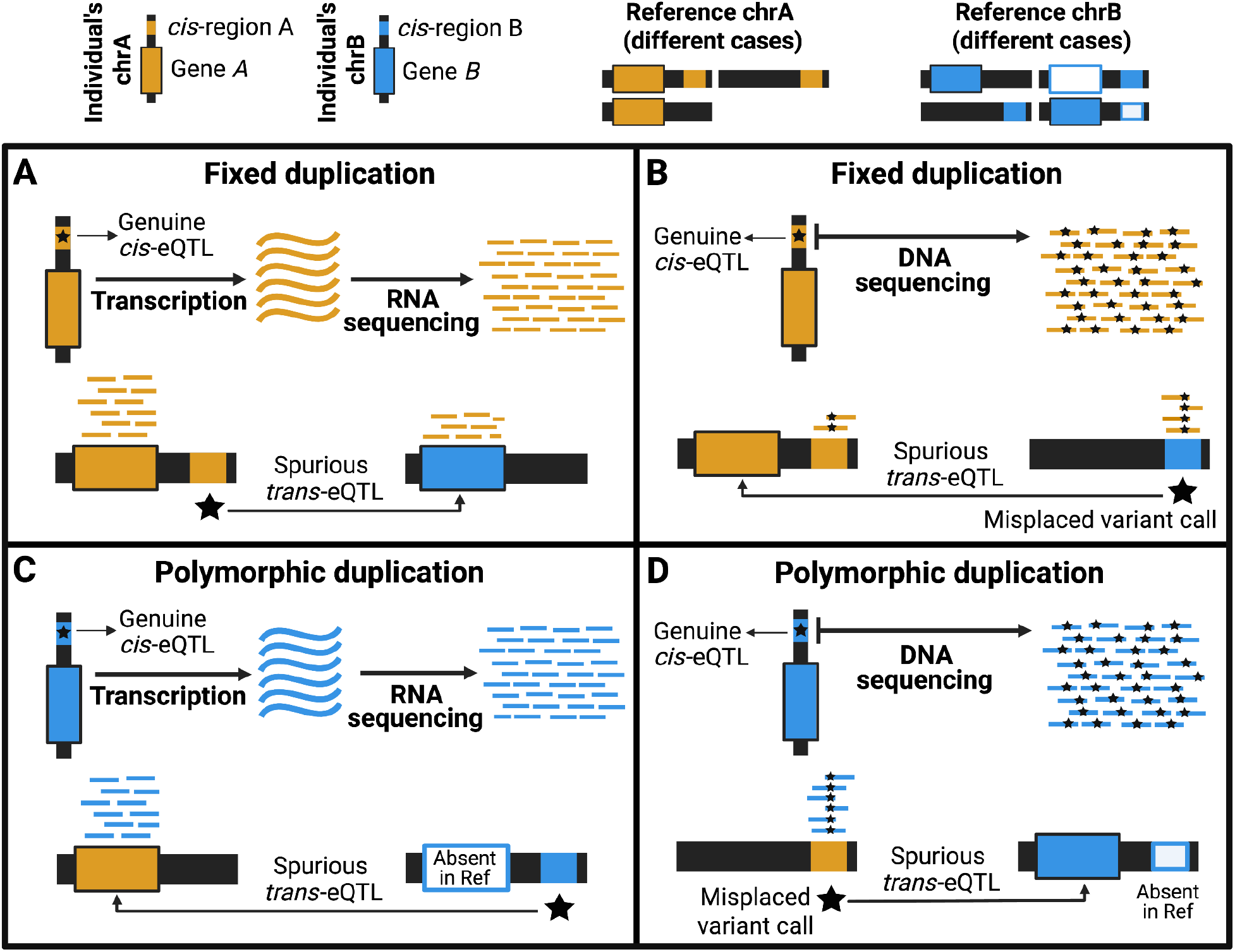
Mismapping of RNA and DNA reads leads to spurious trans-QTL calls. **A**. Category 1: gene duplication in the reference genome. Gene *A* on chrA has a duplicate (gene *B*) on chrB in the reference genome. RNA reads from the transcripts of gene *A* erroneously map onto those of gene *B*, creating a spurious *trans*-association between the genuine *cis*-QTL for gene *A* and the expression level of gene *B*. **B**. Category 2: regulatory-region duplication in the reference genome. The cis-regulatory region for gene *A* on chrA, but not gene *A* itself, is duplicated to chrB in the reference. DNA reads from the cis-regulatory region of gene *A* then map to the duplication region on chrB, leading to the erroneous placement of the *cis*-QTL from chrA to chrB during variant calling. The genuine *cis*-QTL for gene *A* is spuriously assigned as its *trans*-QTL. **C**. Category 3: polymorphic gene duplication absent in the reference genome. Gene *A* on chrA has a polymorphic duplicate (gene *B*) inserted on chrB that is absent from the reference. Both gene copies are transcribed, but RNA reads from the transcripts of gene *B* align to the single reference copy (gene *A*) on chrA. Variants near the insertion site on chrB are in linkage disequilibrium with the duplication and appear as false *trans*-QTLs for gene *A*. **D**. Category 4: polymorphic regulatory-region duplication absent in the reference. A region on chrA is polymorphically duplicated near gene *B* on chrB, but is absent from the reference. The duplicated sequence on chrB carries a *cis*-QTL for gene *B*. Short DNA reads covering that variant map to the single reference copy on chrA. The variant is misplaced to chrA during variant calling. This misplaced variant on chrA appears associated with the expression of gene *B* on chrB, creating a spurious *trans*-QTL.

Saha and Battle recognized and largely resolved category-1 mismapping errors (2018). Their method identifies gene pairs (*A, B*) such that exonic RNA reads from *A* map to the exons of *B* (i.e., “cross-mappable genes”). Then, it removes any putative *trans*-QTL for target gene *A* that lies near the cross-mappable partner gene *B*. This eliminates *cis*-QTLs of gene *A* camouflaged as *trans*-QTLs of gene *B*. The GTEx consortium later incorporated this filter into their QTL pipelines (GTEx Consortium 2020).

However, categories 2–4 of mismapping remain unresolved. To address these, we utilize a simple footprint left by mismapping. Consider two regions, A and B. Region A contains variants in linkage disequilibrium with nearby sites. When reads containing variants from A are mismapped to B, these variants are erroneously called at B. Due to the original linkage disequilibrium pattern, these misplaced variants at B appear correlated with sites near A. Across individuals, this creates an apparent long-range correlation between genotypes at A and B, i.e., genotypic disequilibrium (Weir 1997). It is the genotype analogue of allele-based linkage disequilibrium. Because genotypic disequilibrium does not require haplotype phasing, it can be applied cleanly to distant or interchromosomal window pairs. Notably, this footprint has been used in the past to identify the insertion sites of polymorphic gene duplications absent from the reference (Saitou and Gokcumen 2019). Genuine *trans*-QTLs should not show such genotypic correlations with sites near their target gene; when they do, the *trans*-QTL is likely spurious and should be eliminated.

We applied this framework to GTEx data on *trans*-eQTLs, *trans*-sQTLs, and *trans*-pQTLs (**Figure 2A**). Recall that the GTEx *trans*-QTL pipelines have already resolved category 1 of mismapping. For each *trans*-QTL, defined by a gene-variant-tissue triplet, we carried out the following procedure. First, we subset the GTEx VCF file to include only those individuals that were used to call the QTL in that tissue. We then defined two windows: (i) the target gene coordinates plus 25 kb on each side, and (ii) the *trans*-QTL coordinates plus 25kb on each side. We calculated genotypic disequilibrium (*r*^2^) for all pairs of inter-window variants, and recorded the maximum *r*^2^ as the empirical max(*r*^2^) for that window pair. Next, we generated 200 random size-matched window pairs from the same VCF and repeated the procedure to obtain a null distribution of max(*r*^2^) values. If the empirical max(*r*^2^) exceeded the 95th percentile plus 1.5 times the interquartile range of the null distribution, we inferred that the observed genotypic disequilibrium between the QTL and the gene region was higher than expected without mismapping. When this occurred, we concluded that the *trans*-QTL in question is spurious (**Figure 2B**).

**Figure 2.**
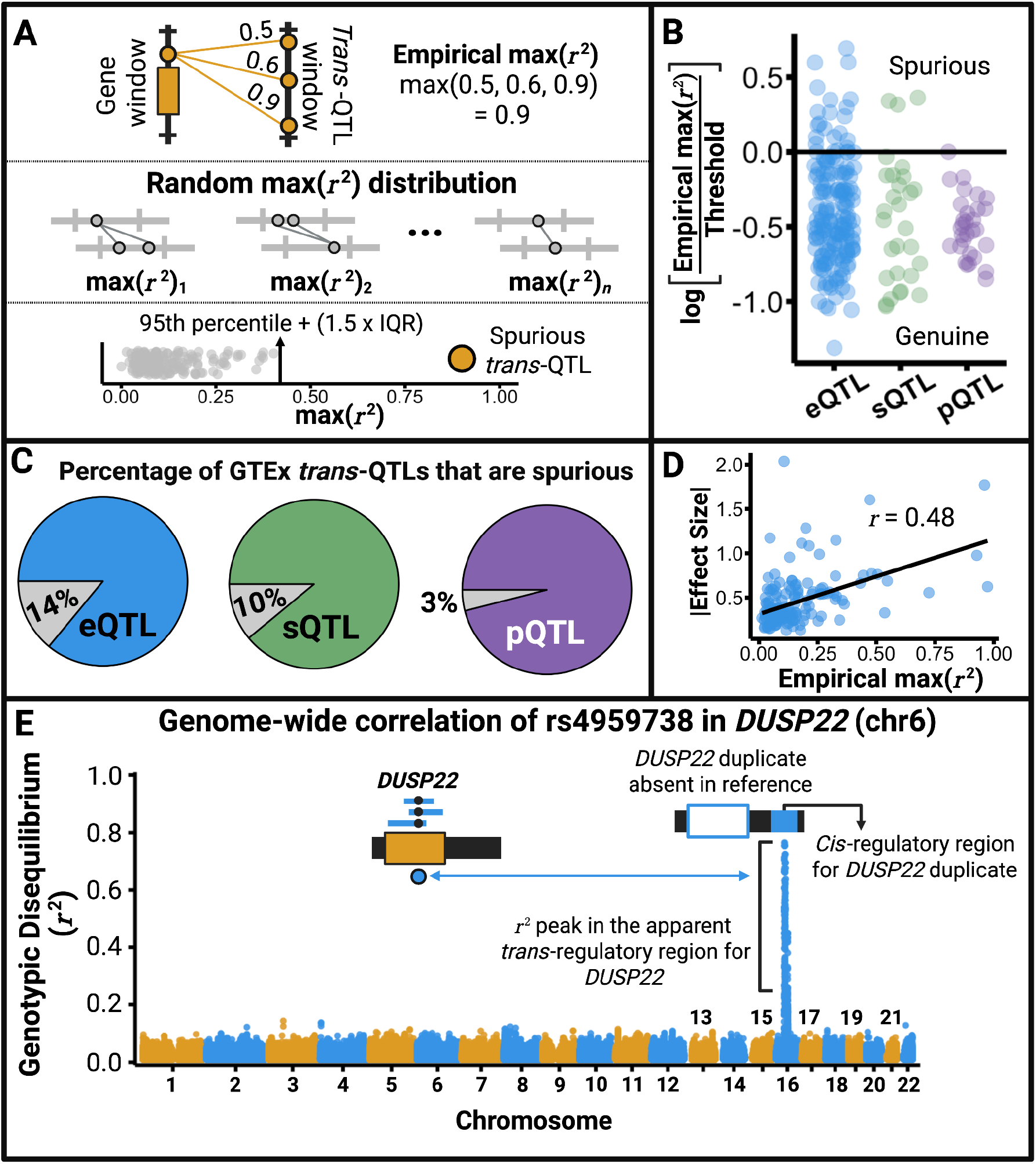
Genotypic disequilibrium between gene and *trans*-QTL windows identifies spurious associations. **A**. Methodological framework. For each *trans*-QTL, we define two windows: the QTL position ±25 kb and the target gene ±25 kb. We calculate genotypic disequilibrium (*r*^2^) across all inter-window variant pairs and record the maximum value, i.e., empirical max(*r*^2^). This empirical max(*r*^2^) is compared to a null distribution from random size-matched window pairs. QTLs with empirical max(*r*^2^) greater than the 95th percentile + 1.5×IQR of the null distribution are flagged as spurious. **B**. Jitter plots of eQTLs, sQTLs, and pQTLs. The y-axis represents the log ratio of empirical max(*r*^2^) to the null threshold (95th percentile + 1.5×IQR). Values >0 indicate spurious QTLs. **C**. Proportion of spurious vs genuine QTLs. Gray represents spurious QTLs; colored represents genuine QTLs. **D**. Correlation between empirical max(*r*^2^) and effect size. **E**. Manhattan plot of genotypic disequilibrium for rs4959738 in *DUSP22*. Based on GTEx individuals included in spleen eQTL analysis, rs4959738 shows strong genotypic disequilibrium with variants in two regions: within the *DUSP22* locus on chromosome 6 and in the apparent spleen *trans*-QTL region on chromosome 16. This long-range disequilibrium reflects a polymorphic duplication of *DUSP22* (category 3, **Figure 1C**), which underlies the spurious *trans* association.

Using this method, we find that 22/161 (14%) *trans*-eQTLs, 3/29 (10%) *trans*-sQTLs, and 1/29 (3%) *trans*-pQTLs reported by the GTEx consortium are spurious (**Figure 2C; Figures S1-3**). These numbers are likely underestimates since we define spurious very conservatively. The pervasiveness of spurious *trans*-QTLs is further illustrated by the positive correlation (r= 0.48) between genotypic disequilibrium and association effect size (**Figure 2D**), consistent with large-effect *cis*-QTLs masquerading as *trans*-QTLs, as noted in the introduction.

As an illustration, the Dual Specificity Phosphatase 22 gene (*DUSP22*) on chromosome 6 shows both a *trans*-eQTL (rs879065486) and a *trans*-sQTL (rs1435502613) on chromosome 16. However, this association is an artifact of genotypic disequilibrium caused by a polymorphic duplication of *DUSP22*, which is inserted on chromosome 16 but absent from the hg38 reference genome (**Figure 1C; Figure 2E**) (Shew et al. 2021). This newer copy of *DUSP22* is present in the more recent Telomere-to-Telomere (T2T) genome (Behera et al. 2023). Several studies have also reported *trans* methylation QTLs on chromosome 16 for CpG sites in *DUSP22* (Boks et al. 2018; Lemire et al. 2015; Bonder et al. 2017; Gaunt et al. 2016), which we posit are likewise artifacts of genotypic disequilibrium arising from this duplication.

## Conclusion

Our analysis shows that a substantial fraction of reported *trans*-QTLs are artifacts of read mismapping stemming from segmental duplications. Such artifacts leave a footprint in the form of long-range genotypic disequilibrium, which we leveraged to distinguish between spurious and genuine *trans*-acting regulatory variants. Our genotypic disequilibrium framework will become especially useful as consortia, such as GTEx, expand their QTL analyses to include multiple developmental stages and non-human primates (Coorens et al. 2025). We caution that it is important to compute genotypic disequilibrium using the same individuals and variant-calling pipelines as those used to generate the associations, to ensure accurate flagging of spurious signals.

We note that long-range genotypic disequilibrium can, in principle, arise from epistatic selection (Kimura 1956). Nevertheless, to explain our results, epistatic selection would have to be unrealistically common, strong, and sustained (*r*^2^ decays by a factor of four each generation without selection). To aid future work on epistatic selection in humans, we have made available the interchromosomal genotypic disequilibrium map for each of the 20 unadmixed populations from the 1000 Genomes Project (**Supplementary data S1**).

The utility of our approach extends beyond *trans*-QTL discovery and is especially relevant given the growing appreciation of segmental duplications across species (Islam et al. 2024). It provides a means of filtering false positives in any analysis that links distant genomic regions or associates multiple genomic regions with phenotypes. In Hi-C data, for example, polymorphic duplications inserted on other chromosomes can generate the illusion of long-distance contacts when reads are mismapped to the reference copy. The reported contacts between *DUSP22* on chromosome 6 and apparent regulatory regions on chromosome 16 (Boks et al. 2018; Rao et al. 2014) are likely such an artifact. Similarly, genome-wide association peaks that appear distant and independent are not truly so if one of them arises from mismapping, since genotypic disequilibrium couples the signals.

Co-expression networks are also susceptible to the mismapping problem. Mismapping of RNA reads can create spurious correlations between genes near the source locus and genes near the mismapped site. Such artifacts can be reduced either by regressing out the effects of genotypic disequilibrium from pairwise correlations or by setting correlations to zero for gene pairs that exceed a genotypic disequilibrium threshold. Approaches such as the one proposed here will become increasingly important as co-expression networks become multi-layered and extended to multiple tissues (Russell et al. 2023), where such errors can accumulate.

Incorporating our framework into these methods will ground inferences about disease, evolution, and gene regulation in true biology rather than technical artifacts.

## Acknowledgements

We thank Kendra Scheer and Dr. Derek J. Taylor for their careful reading of the manuscript. We also thank the Gokcumen lab for their suggestions during the onerous task of choosing an appropriate title for this manuscript. O.G. acknowledges support from the National Science Foundation (under grant nos. 2049947 and 2123284) and the National Institute of Health (R35-GM156519). The funders had no role in study design, data collection and analysis, publication decision, or manuscript preparation.

## Methods

### Data

The Genotype-Tissue Expression (GTEx) project is an ongoing effort to build a comprehensive public resource to study tissue-specific gene expression and regulation. Specifically, we downloaded version 8 *trans*-eQTLs and *trans*-sQTLs (FDR < 0.05) from the GTEx portal (https://gtexportal.org/home/downloads/adult-gtex/qtl). We obtained the *trans*-pQTLs from (Fang et al. 2025). We note that GTEx does not report *trans* methylation QTLs. In all GTEx *trans*-QTL calls, the putative regulatory variant is located on a different chromosome from the target gene. Genomic variant calls were obtained from GTEx protected data via dbGaP accession phs000424.v10.p2 on the anvil platform (https://anvil.terra.bio/#workspaces/anvil-datastorage/AnVIL_GTEx_V9_hg38).

### Interchromosomal genotypic disequilibrium

For each tissue r, we processed the GTEx VCF file (944 individuals) to create the file VCF_*t*_, which contains only the individuals used to call *trans*-QTLs in tissue r. Then, for each *trans*-QTL defined by the gene-variant-tissue triplet (*g,v,t*), we subsetted the VCF_*t*_ to retain variants in two windows: (i) the target gene coordinates ±25 kb, and (ii) the *trans*-QTL coordinates ±25 kb. In all cases, the two windows are located on different chromosomes. We call the resulting file VCF_*g,v,t*_. Genotypic disequilibrium (*r*^2^) was calculated for each inter-window variant pair using VCFtools:

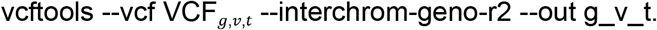

We then calculate the maximum of all *r*^2^ values obtained and call it empirical max(*r*^2^)_*g,v,t*_.

### Null distribution

To investigate whether max(*r*^2^)_*g,v,t*_ was an outlier relative to the expectation under no mismapping, we generated 200 random size-matched interchromosomal window pairs. Before analysis, we masked VCF_*g,v,t*_to remove variants within interspersed repeats, defined as SINEs, LINEs, LTRs, DNA transposons, rolling-circle transposons, or retrotranposons longer than 100 bp. This masking ensured that the null max(*r*^2^) distribution was not inflated by mismapping due to repeats. For each random window pair *i*, we subsetted the masked VCF_*g,v,t*_to retain variants falling within one of the windows in the pair *i*. We call this file VCF_*i*_. Then, we calculated *r*^2^ for all inter-window variant pairs in VCF_*i*_:

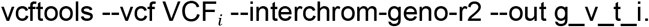

We recorded the maximum *r*^2^ across the inter-window variant pairs and call it max(*r*^2^)_*i*_.

For each, *trans*-QTL (*g,v,t*), we calculated max(*r*^2^)_*i*_for all *i* ∈ {1, …, 200}, yielding a null distribution of max(*r*^2^) values for the *trans*-QTL. If the empirical max(*r*^2^)_*g,v,t*_ exceeded the 95th percentile + 1.5 × interquartile range (IQR) of the null distribution, we inferred that genotypic disequilibrium between the gene window and QTL window was higher than expected under the hypothesis of no read mismapping. Since standard outlier detection uses the 3^rd^ quartile + 1.5 × IQR, our approach is very conservative.

## Supplementary Material

**Figure S1.**
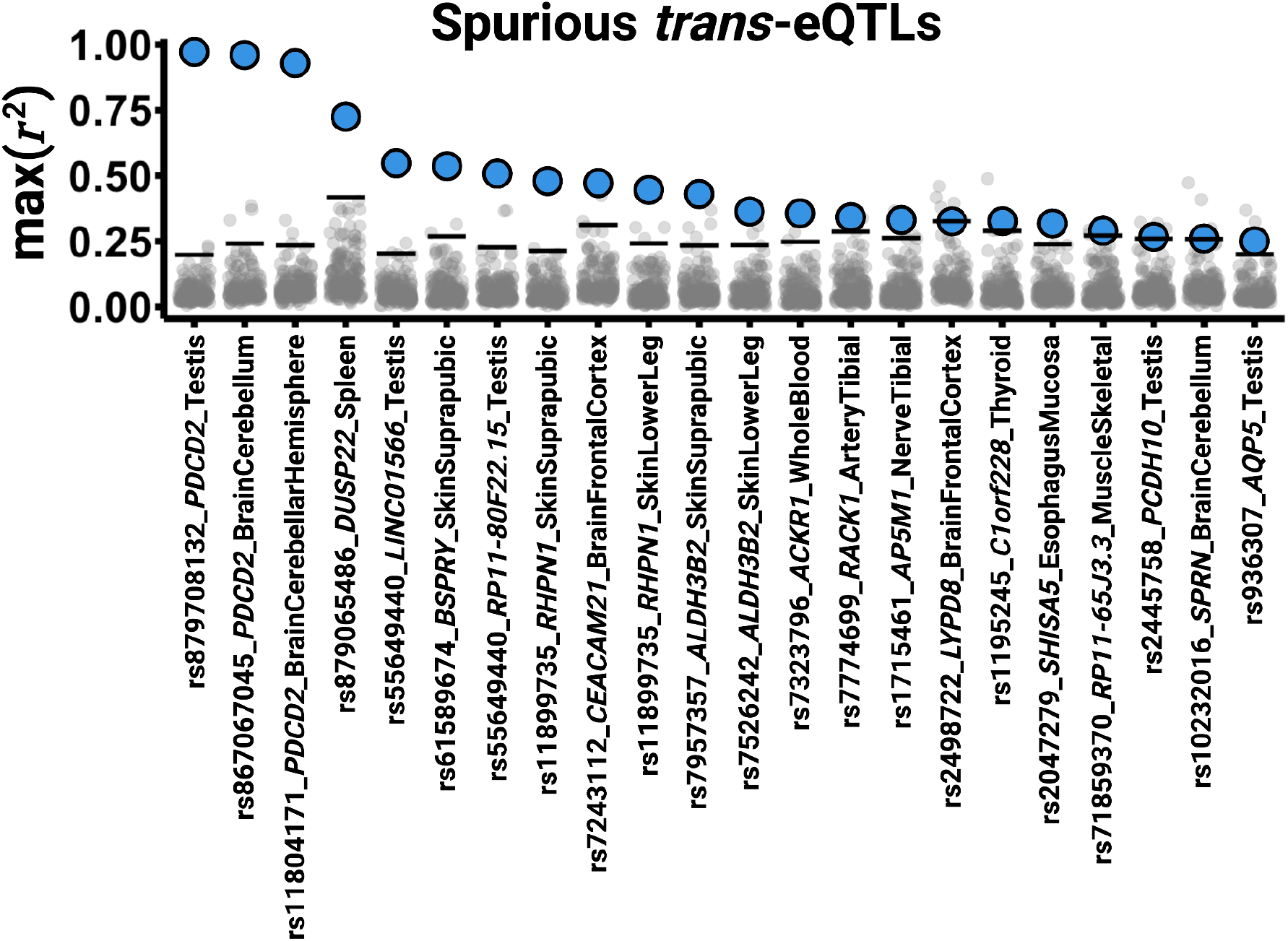
Empirical max(*r*^2^) versus the null distribution of max(*r*^2^) for spurious *trans*-eQTLs.

**Figure S2.**
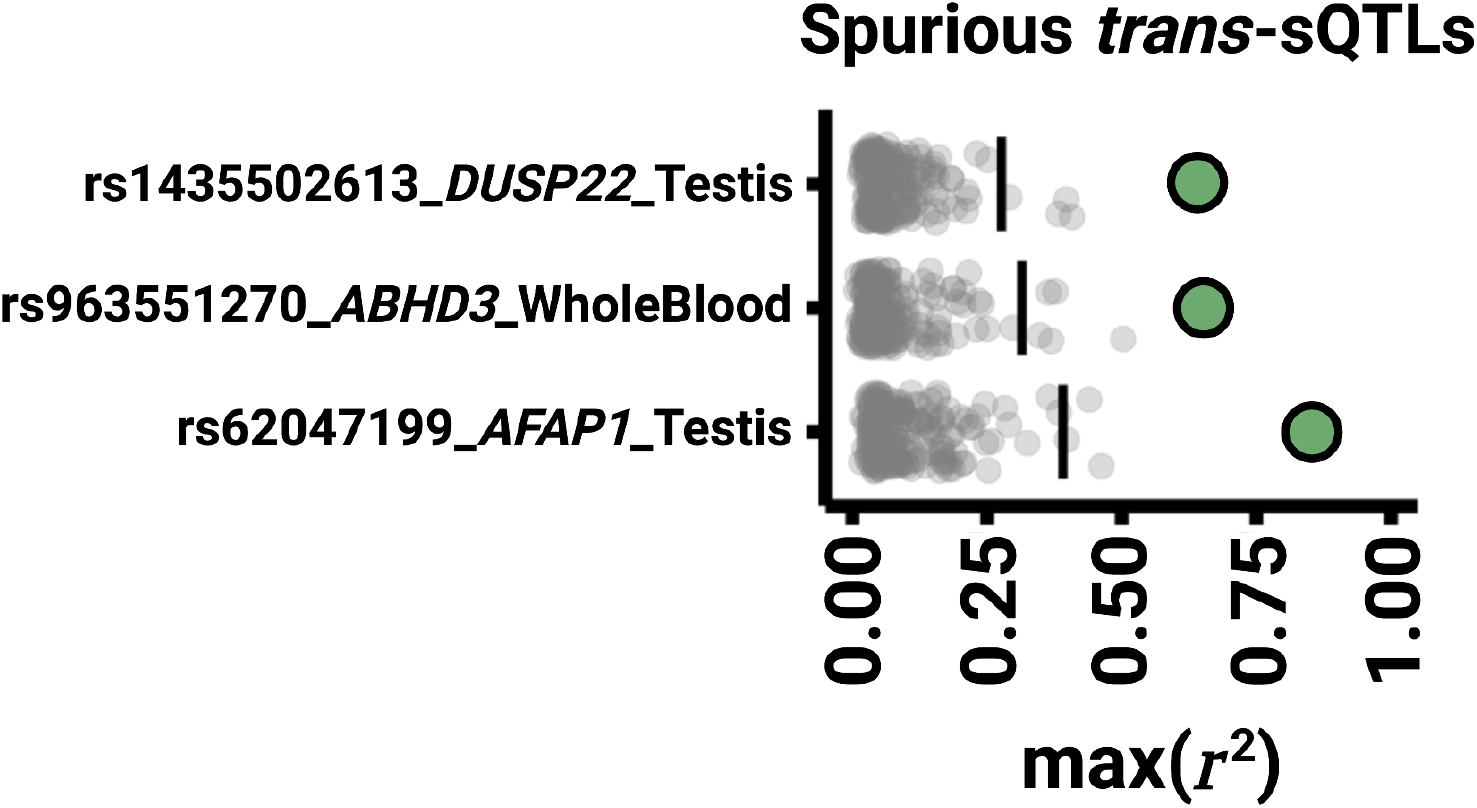
Empirical max(*r*^2^) versus the null distribution of max(r2) for spurious *trans*-sQTLs.

**Figure S3.**
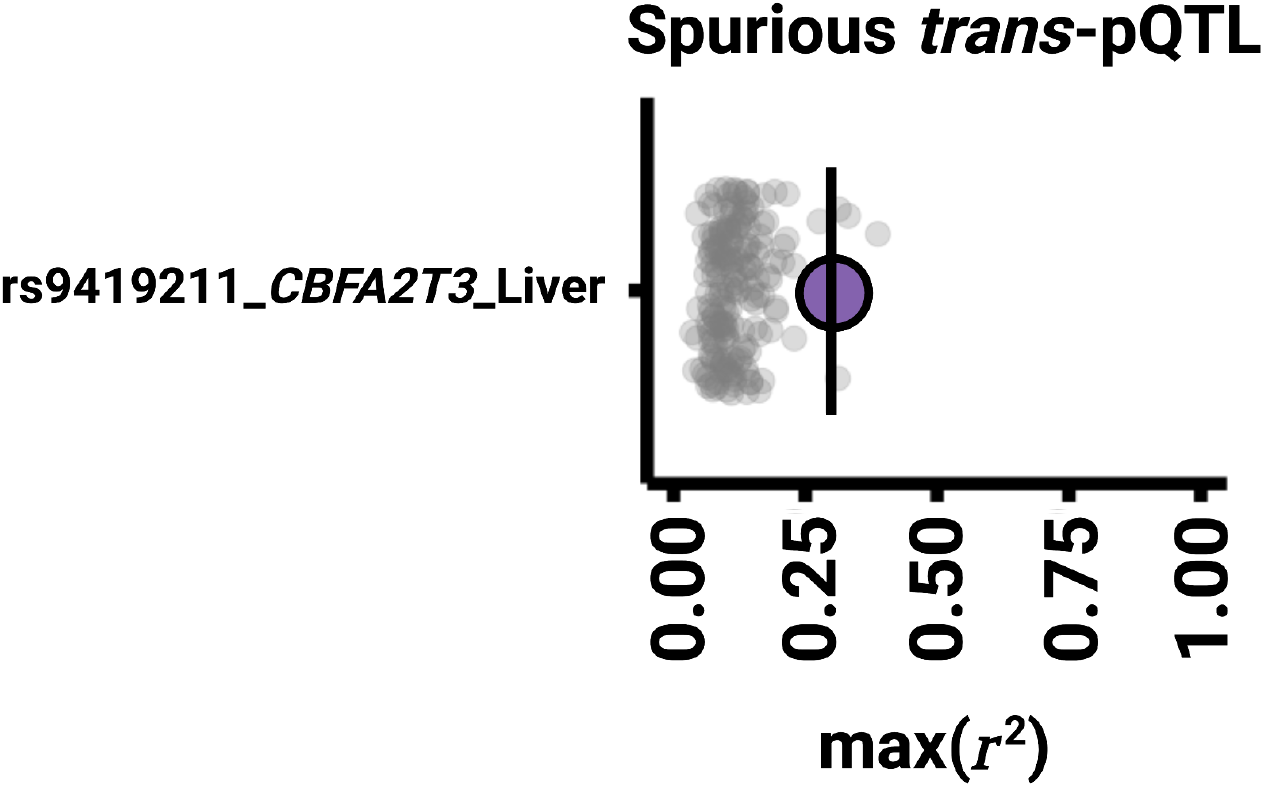
Empirical max(*r*^2^) versus the null distribution of max(*r*^2^) for the spurious *trans*-pQTL.

